# Assessing the Influence of Tractography Methods on Detected White Matter Microstructure in Alzheimers disease

**DOI:** 10.1101/2025.11.11.687747

**Authors:** Yuhan Shuai, Yixue Feng, Julio E. Villalon-Reina, Talia M. Nir, Sophia I. Thomopoulos, Paul M. Thompson, Bramsh Q. Chandio

## Abstract

Tractometry enables detailed mapping of white matter microstructure along individual tracts and is widely used to study disease effects such as those seen in Alzheimer’s disease (AD). However, how different tractography algorithms influence tractometry outcomes remains unclear. Here, we compared whole-brain deterministic and probabilistic tractography using the BUndle ANalytics (BUAN) framework in the Alzheimer’s Disease Neuroimaging Initiative (ADNI) dataset, including 118 AD and 728 cognitively normal (CN) participants. Both approaches revealed the expected pattern of lower fractional anisotropy (FA) and higher mean, radial, and axial diffusivity (MD, RD, AxD) in AD, consistent with white matter degeneration. Despite broadly similar global trends, substantial bundle-level differences emerged between the two tractography methods. Probabilistic tracking produced stronger and more spatially extended effects in the fornix, a small and highly curved limbic pathway vulnerable to AD-related degeneration, whereas deterministic tracking showed greater sensitivity in the posterior segments of the right superior longitudinal fasciculus (SLF R). These discrepancies highlight that the choice of tractography algorithm can alter detecting disease effects, emphasizing the need for cross-method validation to ensure the robustness and interpretability of along-tract measures.

## 1. INTRODUCTION

Alzheimer’s disease is a leading cause of disability and dependency among older adults worldwide, contributing substantially to the global burden of dementia. Diffusion MRI (dMRI) provides a noninvasive means to assess white matter (WM) microstructure by modeling water diffusion in-vivo. Among dMRI models, diffusion tensor imaging (DTI) remains the most widely used framework, offering four primary metrics: fractional anisotropy (FA), mean diffusivity (MD), axial diffusivity (AxD), and radial diffusivity (RD), which reflect the directional coherence and magnitude of water diffusion along axonal pathways. In AD, dMRI studies consistently report reduced FA and elevated diffusivity metrics compared to cognitive normal individuals [1, 2, 3]. These changes indicate loss of axonal integrity and myelin disruption, consistent with the progressive neural disconnection and WM degeneration observed in AD [4, 5]. Such microstructural alterations often appear early in the disease course and are closely associated with both pathological hallmarks and cognitive decline [6].

Tractometry quantifies microstructure along reconstructed WM pathways derived from tractography, and has been used to study AD [7, 8] and the effects of its hallmark pathologies, which include abnormal accumulation of brain amyloid and tau [9]. However, different tractography algorithms may result in inconsistent estimates of the spatial distribution and magnitude of WM alterations [10, 11]. Deterministic tractography, for example, propagates streamlines along the principal diffusion direction estimated locally from the dMRI data. As a result, streamlines originating at a given seed point will always follow the same trajectory, producing consistent paths that represent the dominant orientation [12, 13]. In contrast, probabilistic tractography samples multiple orientations at each step, and accounts for uncertainty in local fiber orientations, allowing for greater exploration of alternative pathways and better coverage. However, this sampling approach can also introduce variability in tract reconstruction [14, 15]. In principle, probabilistic approaches, by incorporating greater directional uncertainty, can typically achieve broader bundle coverage, that is, a more extensive sampling of voxels potentially belonging to a tract, capturing peripheral or branching fibers that may be missed by deterministic algorithms [16]. Alternatively, deterministic methods often produce more spatially restricted pathways with fewer false positives [17], and may be especially valuable where high precision is crucial at the individual level, such as in surgical planning [2, 18]. These fundamental differences raise a critical question: do tractography methods systematically influence the outcomes of tractometry studies of brain diseases?

In this study, we analyzed data from the Alzheimer’s Disease Neuroimaging Initiative to compare how WM bundles reconstructed from deterministic vs. probabilistic tractography affect along-tract analysis of FA, MD, AxD, and RD measures using the BUAN tractometry framework [19]. We set out to assess whether these methods would produce different effect sizes or statistical significance patterns for bundle-wise differences in AD-related abnormalities. By quantifying algorithm-induced variability, we aimed to determine whether tractography approaches differ in their overall sensitivity to AD-related microstructural alterations, and in the spatial profile of abnormalities detected across white matter bundles.

## 2. METHODS

### 2.1. Diffusion MRI Processing

We analyzed diffusion MRI data from the ADNI3 cohort, including cognitively normal (CN, *N* = 728) and Alzheimer’s disease (AD, *N* = 118) participants. The mean age was 71.3 *±* 7.9 years for CN. Females represented 62.2% of the CN group (453/728) and 45.8% of the AD group (54/118). Diffusion scans were collected across multiple ADNI sites using seven imaging protocols (denoted by S55, GE54, S127, S31, P33, P36, and GE36).^1^ All standard preprocessing steps were performed following the established ADNI dMRI pipeline [3], denoising, including motion and eddy current correction, brain extraction, and diffusion tensor fitting. Diffusion data were resampled to an isotropic resolution of 1.5 *×* 1.5 *×* 1.5 mm^3^. Whole-brain tractograms were generated using constrained spherical deconvolution (CSD) and both local deterministic and probabilistic tracking for each subject using DIPY [20]. For both tracking methods, the following parameters were held identical: tracking was seeded from voxels with FA *>* 0.15, using eight seeds per voxel, and streamlines were propagated with a step size of 0.5 mm with a minimum separation angle of 25° was set. Tracking was terminated when FA dropped below 0.15.

Whole-brain tractograms were affinely registered to an atlas in the MNI space using the streamline-based linear registration method [20]. Fourty-two WM bundles were then extracted from the registered tractograms using an auto-calibrated implementation of RecoBundles [21, 19], with the HCP842 atlas bundles [22] used as reference models for streamline segmentation. The same parameter settings were applied for both deterministic and probabilistic tractograms. As a quality control (QC) step, we evaluated the spatial correspondence between each individual bundle and the group average density map using the normalized cross-correlation (NCC) metric [23]. For each bundle, participants whose NCC values deviated by more than 3 standard deviations from the group mean were excluded from subsequent analysis. Reconstruction failed for more than 70% of subjects for the anterior and posterior commissure, so these regions were also excluded from the analysis.

### 2.2. BUAN Tractometry

Each extracted bundle was divided into 5 mm segments along its trajectory using the BUAN framework [19]. Model bundle centroids were defined in a common template space and clustered with QuickBundles to generate reference centroid points along each tract. For every subject, each streamline point was assigned to its nearest centroid based on Euclidean distance, producing segment-wise assignment maps for subsequent analyses. Microstructural measures were mapped onto the tracts in native space.

Segment-wise associations were assessed using linear mixed-effects (LME) models of the form:

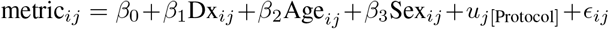

where *metric* represents each diffusion measure, *Dx* represents diagnostic group (CN or AD), and *Protocol* was included as a random intercept to account for variability in dMRI acquisitions. Age and sex were entered as fixed covariates. A global Benjamini–Hochberg false discovery rate (FDR) correction was applied to adjust for multiple comparisons across all segments across all bundles[24]. For each segment, the partial effect size *d* was calculated with reference to the residual standard deviation. To compare our findings, we calculated the *global FDR proportion*, defined as the number of bundle segments that passed FDR relative to the total number of segments tested across all 40 bundles; each segment was counted once, without weighting by bundle length or the number of streamlines. We also calculated the mean partial *d* across all segments from all bundles and the mean partial *d* for the segments with significant results. The Dice overlap between significant segments across two methods where computed to evaluate the similarity of AD patterns identified by our tractometry analysis.

## 3. RESULTS

The FDR proportion, mean partial *d* are shown in **Table 1** Across all diffusion metrics, deterministic and probabilistic tractography produced comparable results overall (Table 1; Fig. 1). Deterministic tractography shows higher FDR proportion by roughly 0.03, suggesting slightly higher overall sensitivity to AD-related effects. Probabilistic tractography shows marginally higher mean effect sizes (partial *d*) across all segments and significant segments for diffusivity measures but not for FA. Dice exceeded 0.90 for all metrics: MD patterns were the most similar and FA the least. Across both methods, MD exhibited the most extensive and significant diagnostic effects, followed by RD, AxD, and FA. Overall, the spatial extent and direction of effects were broadly comparable between methods, indicating reproducibility of disease-related WM alterations, in this case.

**Table 1.** Comparison of deterministic and probabilistic tractography across diffusion metrics. In each case, the method with a numerically greater effect is shown in **bold**.

|  | Metric | AxD | FA | MD | RD |
| --- | --- | --- | --- | --- | --- |
| FDR proportion | det | **0.493** | **0.325** | **0.660** | **0.583** |
|  | prob | 0.470 | 0.283 | 0.627 | 0.558 |
| Mean $d$ (all) | det | 0.266 | **-0.149** | 0.367 | 0.323 |
|  | prob | **0.272** | -0.143 | **0.369** | 0.323 |
| Mean $d$ (significant) | det | 0.438 | **-0.284** | 0.509 | 0.502 |
|  | prob | **0.442** | -0.278 | **0.512** | **0.503** |
| Dice |  | 0.938 | 0.906 | 0.955 | 0.948 |

**Fig. 1.**
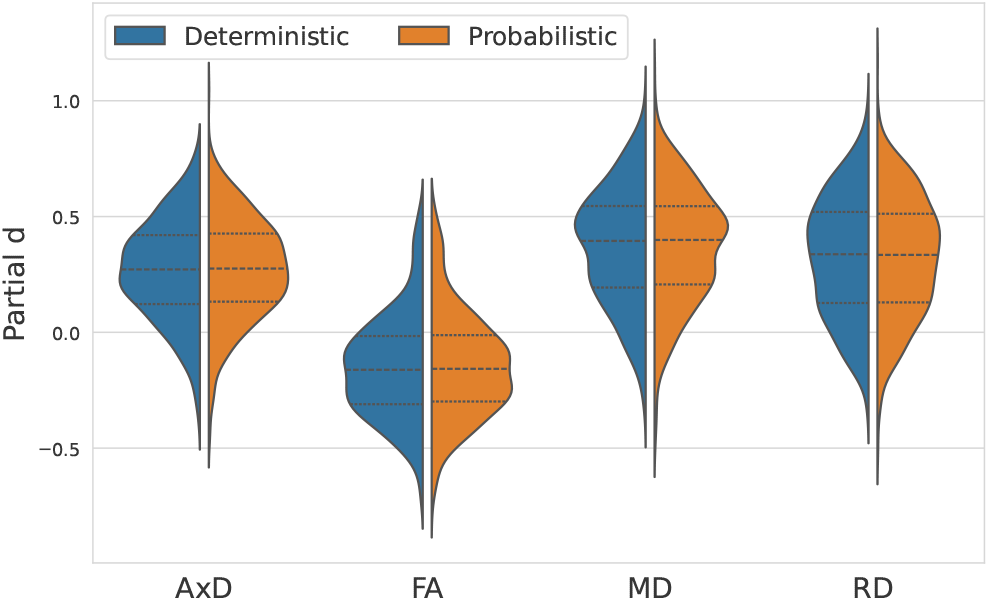
Distribution of segment-wise effect sizes (partial d) across diffusion metrics for deterministic and probabilistic tractography methods. Both methods show similar effect-size distributions, indicating comparable sensitivity across metrics, in this case.

Among 40 WM bundles examined, the most extensive AD effects were found in the bilaterial inferior longitudinal fasciculus (ILF) and the corpus callosum forceps major (CC ForcepsMajor), where both tracking methods identified significant results in more than 95% of these bundles for all diffusivity measures. Probabilistic tractography produced higher mean effect size in diffusivity measures for the CC ForcepsMajor but lower mean effect size in ILF L, though the difference in effect size was no greater than 0.05.

To identify bundles with the greatest variability across methods, we calculated, for each tract and diffusion metric, the difference in FDR proportion and the absolute difference in mean effect size (partial *d*) between deterministic and probabilistic tractography. The left fornix showed the largest difference in effect size across methods and the right superior longitudinal fasciculus (SLF R) showed the greatest difference in FDR proportion. The along-tract profiles for four DTI metrics from both tractography methods are shown in (**Fig. 2**). For F L, deterministic tractography did not identify any significant results for diffusivity measures, and probabilistic tractography produced higher effect size on average across segments, by 0.431 for MD, 0.42 for AxD and 0.416 for RD. For FA, however, probabilistic tractometry did not identify significant results, and had lower mean effect size by 0.097. We identified different patterns for the SLF R, where deterministic tracking yielded more significant segments than probabilistic tractography for diffusivity measures, with the difference in bundle FDR proportion being 0.278 for AxD, and 0.222 for MD and RD. Both methods produced identical significant segments in FA. Deterministic tractography also produced higher along-tract effect size in the posterior portion of SLF R.

**Fig. 2.**
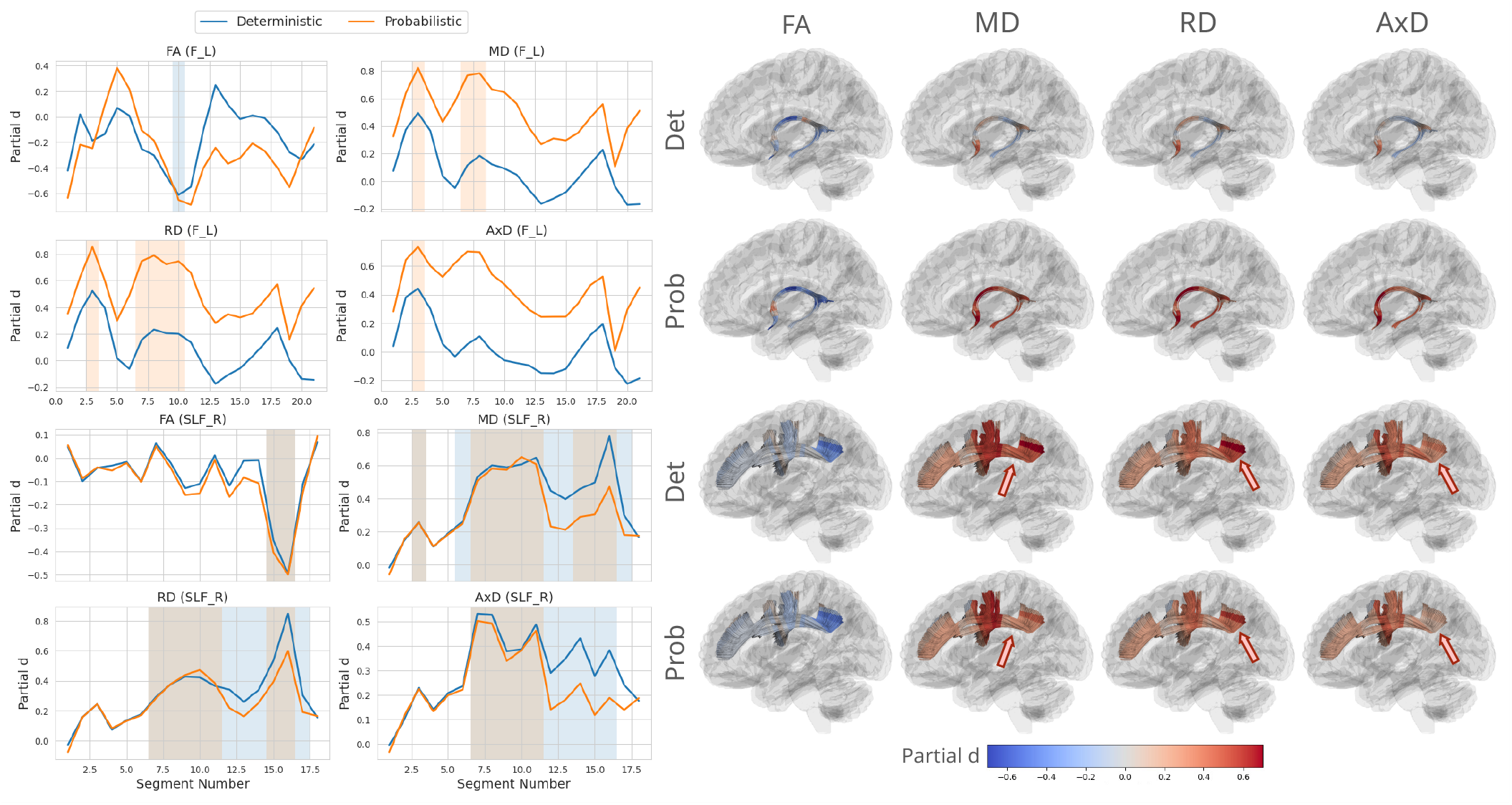
Comparison of along-tract profiles and spatial effect maps between deterministic and probabilistic tractography. ***Left:*** Segment-wise partial *d* values for diffusion metrics (FA, MD, RD, AxD) plotted along the tract for the left fornix (F L) and right superior longitudinal fasciculus (SLF R). Shaded regions indicate tract segments showing statistically significant group differences after global FDR correction (q *<* 0.05). ***Right:*** 3D renderings visualize the spatial distribution of partial *d* values along each tract (blue = negative, red = positive). Probabilistic tracking shows stronger effects in the fornix, while deterministic tracking highlights posterior SLF effects, illustrating tract-specific variability between methods.

## 4. DISCUSSION

In this study, we systematically compared deterministic and probabilistic tractography using ADNI diffusion MRI data to examine how this algorithmic choice influences along-tract diffusion metrics in detecting AD related group effects using tractometry. We found highly consistent global patterns across tractography methods, demonstrating that along-tract tractometry results remain stable overall despite differences in reconstruction algorithms. Both tractography methods consistently identified microstructural alterations in DTI metrics, with overall reduced FA and increased diffusivity.

Despite the global agreement, localized discrepancies emerged in several bundles, reflecting algorithm-dependent effects. Of the bundles examined, the left fornix (F_L) and right superior longitudinal fasciculus (SLF_R) showed the largest differences across tractography methods, where probabilistic tracking produced stronger and more spatially extended effects in the fornix, and deterministic tracking showed greater sensitivity in the posterior SLF. The fornix, a small and highly curved limbic pathway, is challenging to reconstruct, yet it is of interest in dementia studies, as it is particularly vulnerable to AD-related degeneration [25]. As the fornix serves as the principal output of the hippocampus within the limbic memory circuit, it is among the earliest WM pathways to show demyelination and atrophy in AD. Subtle reconstruction differences may therefore interact with the true disease effects [26]. In contrast, the SLF R, a long association tract linking frontal and parietotemporal regions, showed slightly higher sensitivity under deterministic tracking, likely reflecting its extended and branching geometry that probabilistic tracking may represent with greater angular dispersion [27]. These tract-specific effects underscore how local anatomy and disease vulnerability can affect algorithmic outcomes, even when global results remain generally consistent across methods.

Several limitations should be noted. First, our comparison was conducted within a single tractography and tractometry ecosystem using matched parameters; therefore, the observed differences may depend on specific pipeline implementations. Although both tractography methods were implemented with standardized settings, alternative processing pipelines or parameter choices, such as seeding density, stopping criteria, or streamline filtering thresholds, could further influence bundle reconstruction and along-tract estimates. Second, uncertainty estimates for effect-size differences (e.g., via bootstrapping) were not computed and should be incorporated in future work, particularly in studies with smaller sample sizes or weaker effects. In addition to automated quality control using quantitative metrics [23], manual inspection of challenging bundles or those with high inter-subject variability may further elucidate algorithm-dependent differences. Finally, while Alzheimer’s disease is associated with relatively large white matter effects, algorithmic choices may play a more critical role when effects are subtler, such as in psychiatric populations or borderline-powered studies. Future work will examine whether tract shape differences across tractography methods are consistent across datasets and acquisition protocols.

## 5. ACKNOWLEDGMENTS

This work was supported by AARG-23-1149996, R01AG058854, R01AG087513, RF1AG057892, U01AG068057, U19AG024904, S10OD032285.

## Footnotes

1 The abbreviations here denote whether Siemens (S), General Electric (GE), or Philips scanners were used, and the number denotes the total number of volumes acquired.

## REFERENCES

[1] P. J. Winklewski et al., “Understanding the physiopathology behind axial and radial diffusivity changes— what do we know?,” Frontiers in Neurology, vol. 9, pp. 92, 2018.

[2] Artemis Zavaliangos-Petropulu et al., “Diffusion MRI indices and their relation to cognitive impairment in brain aging: The updated multi-protocol approach in Alzheimer’s Disease Neuroimaging Initiative (ADNI3),” Frontiers in Neuroinformatics, vol. 13, no. 2, 2019.

[3] Sophia I. Thomopoulos et al., “Diffusion MRI metrics and their relation to dementia severity: effects of harmonization approaches,” in Proceedings of SPIE – The International Society for Optical Engineering, 17th International Symposium on Medical Information Processing and Analysis, Dec. 2021, vol. 12088, p. 120880K.

[4] Inge K. Amlien and Anders M. Fjell, “Diffusion tensor imaging of white matter degeneration in Alzheimer’s disease and mild cognitive impairment,” Neuroscience, vol. 276, pp. 206–215, 2014.

[5] T. M. Nir et al., “Effectiveness of regional DTI measures in distinguishing Alzheimer’s disease, MCI, and normal aging,” NeuroImage: Clinical, vol. 3, pp. 180–195, 2013.

[6] B. T. Gold et al., “White matter integrity and vulnerability to Alzheimer’s disease: Preliminary findings and future directions,” Biochimica et Biophysica Acta (BBA) - Molecular Basis of Disease, vol. 1822, no. 3, pp. 416–422, 2012.

[7] Yixue Feng et al., “Microstructural mapping of neural pathways in Alzheimer’s disease using macrostructure-informed normative tractometry,” Alzheimer’s & Dementia, p. alz.14371, Dec. 2024.

[8] Bramsh Q Chandio et al., “Microstructural changes in the white matter tracts of the brain due to mild cognitive impairment,” Alzheimer’s & Dementia, vol. 18, pp. e065339, 2022.

[9] Bramsh Qamar Chandio et al., “Amyloid, tau, and APOE in Alzheimer’s Disease: Impact on white matter tracts,” Pacific Symposium on Biocomputing (PSB), 2025.

[10] Klaus H. Maier-Hein et al., “The challenge of mapping the human connectome based on diffusion tractography,” Nature Communications, vol. 8, no. 1, pp. 1349, 2017.

[11] X. Zhang et al., “Characterization of white matter changes along fibers by automated fiber quantification in the early stages of Alzheimer’s disease,” NeuroImage: Clinical, vol. 22, pp. 101723, 2019.

[12] T. E. Conturo et al., “Tracking neuronal fiber pathways in the living human brain,” Proceedings of the National Academy of Sciences, vol. 96, pp. 10422–10427, 1999.

[13] S. Mori et al., “Three-dimensional tracking of axonal projections in the brain by magnetic resonance imaging,” Annals of Neurology, vol. 45, no. 2, pp. 265–269, 1999, Official Journal of the American Neurological Association and the Child Neurology Society.

[14] G. J. M. Parker et al., “A framework for a streamline-based probabilistic index of connectivity (PICo) using a structural interpretation of MRI diffusion measurements,” Journal of Magnetic Resonance Imaging, vol. 18, no. 2, pp. 242–254, 2003.

[15] T. E. J. Behrens et al., “Characterization and propagation of uncertainty in diffusion-weighted MR imaging,” Magnetic Resonance in Medicine, vol. 50, no. 5, pp. 1077–1088, 2003.

[16] Saad Jbabdi and Heidi Johansen-Berg, “Tractography: Where do we go from here?,” Brain Connectivity, vol. 1, no. 3, pp. 169–183, 2011.

[17] Y. Shuai et al., “Deterministic versus probabilistic tractography: Impact on white matter bundle shape,” bioRxiv, 2025, Preprint, bioRxiv: the preprint server for biology.

[18] Fang-Cheng Yeh, “DSI studio: an integrated tractography platform and fiber data hub for accelerating brain research,” Nature Methods, vol. 22, no. 8, pp. 1617–1619, 2025.

[19] Bramsh Qamar Chandio et al., “Bundle analytics, a computational framework for investigating the shapes and profiles of brain pathways across populations,” Scientific Reports, vol. 10, no. 1, pp. 17149, 2020.

[20] Eleftherios Garyfallidis et al., “Robust and efficient linear registration of white-matter fascicles in the space of streamlines,” NeuroImage, vol. 117, pp. 124–140, 2015.

[21] Eleftherios Garyfallidis et al., “Recognition of white matter bundles using local and global streamline-based registration and clustering,” NeuroImage, vol. 170, pp. 283–295, 2018.

[22] Fang-Cheng Yeh et al., “Population-averaged atlas of the macroscale human structural connectome and its network topology,” NeuroImage, vol. 178, pp. 57–68, 2018.

[23] Yixue Feng and others., “Streamline Density Normalization: A Robust Approach to Mitigate Bundle Variability in Multi-Site Diffusion MRI,” Aug. 2025.

[24] Yoav Benjamini and Yosef Hochberg, “Controlling the false discovery rate: A practical and powerful approach to multiple testing,” Journal of the Royal Statistical Society: Series B (Methodological), vol. 57, no. 1, pp. 289–300, 1995.

[25] François Rheault et al., “Bundle-specific fornix reconstruction for dual-tracer PET-tractometry,” bioRxiv, 2018, preprint.

[26] Michelle M. Mielke et al., “Fornix integrity and hippocampal volume predict memory decline and progression to Alzheimer’s disease,” Alzheimer’s & Dementia, vol. 8, no. 2, pp. 105–113, 2012.

[27] A. Kamali et al., “Tracing superior longitudinal fasciculus connectivity in the human brain using high resolution diffusion tensor tractography,” Brain Structure and Function, vol. 219, no. 1, pp. 269–281, 2014.

